# Polyamine dysregulation converges with RASopathies on RAS/MAPK and sensory processing phenotypes in *Drosophila*

**DOI:** 10.64898/2026.01.23.701032

**Authors:** Boyd van Reijmersdal, Melanie de Wit, Kai Prins, Zubi Aimee Govers, Pleuni Schreurs, Marina Boon, Marije Been, Spencer G. Jones, Annette Schenck

## Abstract

RASopathies are developmental conditions associated with cognitive and sensory processing impairments. They are caused by pathogenic variants in genes that result in overactivation of the RAS/MAPK signaling pathway. Genes linked to this pathway have been reported to be enriched among *Drosophila* models with habituation deficits, a behavioral phenotype reflecting sensory filtering. To identify hidden RASopathies – monogenic disorders that converge on RAS/MAPK overactivation without being classically linked to the pathway – we generated 89 and screened 41 viable habituation-deficient *Drosophila* RNAi models for RAS/MAPK overactivation, measured as an increased phosphorylated ERK to ERK ratio. This screen identified *Sms*, the ortholog of human spermine synthase (*SMS*), implicated in Snyder-Robinson syndrome. RAS/MAPK overactivation along with hyperreactivity and habituation impairments are confirmed in a full loss-of-function mutant. A RNAi screen targeting polyamine pathway genes identified *Sat* (human *SAT1/2*, *SATL1*) to reproduce these phenotypes. Knockdown of *Sms* or *Sat* in GABAergic neurons impaired habituation, implicating polyamine metabolism in inhibitory circuit function. These findings reveal previously unrecognized convergence between polyamine dysregulation and RASopathies, suggesting shared therapeutic opportunities through modulation of either pathway.

**SUMMARY STATEMENT:** Using *Drosophila*, we uncovered polyamine metabolism genes, *Sms* and *Sat*, as modulators of RAS/MAPK and sensory processing, revealing a shared mechanism between polyaminopathies and RASopathies that may inform unified therapies.

## INTRODUCTION

RASopathies are a group of developmental conditions caused by pathogenic variants in genes encoding components of the RAS/mitogen-activated protein kinase (RAS/MAPK) signaling pathway, which plays a key role in cell growth, differentiation, and development. These include Neurofibromatosis type 1 (NF1, OMIM #162200), Noonan syndrome (OMIM #163950) and Costello syndrome (OMIM #218040), among others, and are characterized by a range of clinical features including intellectual disability (ID), autism spectrum disorder (ASD), musculoskeletal abnormalities, and increased tumor risk (Alfieri et al., 2014; Rauen, 2013; Rauen, 2022; Tartaglia et al., 2022; Zenker, 2022). Beyond its well-established role in oncogenesis, dysregulation of the RAS/MAPK pathway has been robustly linked to cognitive (dys)function. Recent studies suggest that pharmacological inhibition of this pathway using MEK inhibitors, small-molecule compounds that block MEK, a kinase downstream of RAS, can lead to improvements in cognitive performance in individuals with NF1 (Lalancette et al., 2024; Walsh et al., 2021). These findings suggest that cognitive impairments resulting from dysregulated RAS/MAPK signaling may be amenable to treatment through pharmacological modulation of this pathway, highlighting its promise as a therapeutic target.

Habituation is one of the most fundamental and evolutionarily conserved forms of learning, enabling organisms to suppress responses to repetitive, irrelevant stimuli. By supporting attentional filtering, habituation prevents sensory overload and allows cognitive resources to focus on salient inputs (McDiarmid et al., 2017). Deficits in habituation have been reported in individuals with NF1 and SYNGAP1-associated RASopathies and their animal models (Carreno-Munoz et al., 2021; Pride et al., 2023; Wolman et al., 2014). *Drosophila melanogaster* provides a powerful genetic model to investigate the molecular basis of habituation learning and its disruption in neurodevelopmental disorders (NDDs) (Blok et al., 2022). The light-off startle paradigm, which measures jump responses to repeated light-off stimuli, enables efficient, quantitative assessment of habituation, and has been shown to capture cognitive phenotypes across multiple fly models of ID and ASD. A previous large-scale screen identified 98 *Drosophila* orthologs of genes associated with ID with robust habituation deficits upon pan-neuronal knockdown (Fenckova et al., 2019). Among these, genes operating in RAS/MAPK signaling emerged centrally, and manipulations resulting in increased RAS/MAPK signaling in GABAergic neurons alone was sufficient to impair habituation. These findings highlight habituation learning as a sensitive functional readout to uncover convergent disease mechanisms and potentially treatments that are applicable across genetically diverse NDDs, including those involving RAS/MAPK signaling.

Building on this foundation, we screened previously published and unpublished ID-associated genes that were linked to habituation deficits in *Drosophila* using our light-off startle paradigm to identify genes not previously linked to RAS/MAPK signaling that may converge on this pathway. This approach revealed previously unrecognized RAS/MAPK dysregulation upon loss of several genes, including spermine synthase (*Sms*), associated with Snyder-Robinson syndrome (SRS, OMIM #309583) (Cason et al., 2003; Snyder and Robinson, 1969). Manipulation of additional polyamine pathway genes revealed further mechanistic connections to RAS/MAPK signaling, specifically in the shared context of disrupted sensory processing. This convergence underscores how genetically distinct rare disorders may share common, druggable signaling disruptions and supports the development of unified therapeutic strategies for both polyaminopathies and RASopathies, for which treatment options at present remain limited.

## RESULTS

### A screen of habituation-deficient NDD models identifies RAS/MAPK overactivation upon *Sms* knockdown

To uncover NDD genes that converge on RAS/MAPK pathway activity, we focused on 89 *Drosophila* models (**Table S1**) previously shown to exhibit habituation deficits following pan-neuronal RNAi-mediated knockdown. These genes were drawn from a larger pool (**Table S2**, see Materials and Methods) of over 100 habituation-deficient lines identified in previously published and unpublished high-throughput behavioral screens (Fenckova et al., 2019; Stessman et al., 2017) and single gene studies (Castells-Nobau et al., 2019; De Hayr et al., 2025; Dias et al., 2022). Given that increased RAS/MAPK signaling is sufficient to impair habituation in flies (Fenckova et al., 2019), we hypothesized that elevated pathway activity might underlie the behavioral phenotypes in a subset of these models. To test this hypothesis, we quantified phosphorylated ERK (pERK) and ERK levels via Enzyme-Linked Immunosorbent Assay (ELISA) and used increased ratios of pERK to ERK (pERK/ERK) as a proxy for RAS/MAPK pathway overactivation characterizing RASopathies. To avoid dilution by non-manipulated tissues, we knocked down each gene ubiquitously by crossing an *Act-Gal4* driver to the respective UAS-RNAi and background control lines and measured pERK and ERK levels in adult fly whole head lysates.

Of the 89 RNAi models, 41 were viable to adulthood and thus further investigated (**Fig. 1A**). We identified eight models with increases in pERK/ERK ratios exceeding a 1.2-fold threshold, three of which had an FDR < 0.05 (**Fig. 1B, Table S1**). Two of these three are *Nf1* (2.2-fold) and *Spred* (1.6-fold), which encode established negative regulators of RAS and whose human orthologues are causally implicated in the classic RASopathies NF1 and Legius syndrome (OMIM #611431), respectively. Their identification provides proof of principle that our screen can detect biologically meaningful RAS/MAPK overactivation. The third significant hit was *Sms* (1.7-fold), which encodes the enzyme spermine synthase. In humans, pathogenic variants in *SMS* cause Snyder-Robinson syndrome, an X-linked NDD characterized by ID, muscle hypotonia, and skeletal abnormalities (Cason et al., 2003; Snyder and Robinson, 1969). A significant but <1.2-fold increase was observed for *CG12118* (*MMADHC*), and additional genes displayed non-significant increases >1.2-fold, including *Set2* (*SETD2*), *Nmdar2* (*GRIN2A/B*), *β-Man* (*MANBA*), *ScpX* (*SCP2*), and *dnc* (*PDE4A-D*). In contrast to these modest effects, the significant elevation in *Sms*, a gene not previously molecularly linked to RAS/MAPK signaling yet recovered alongside the well-established RAS/MAPK regulators *Nf1* and *Spred*, revealed a potentially novel pathway connection.

**Figure 1.**
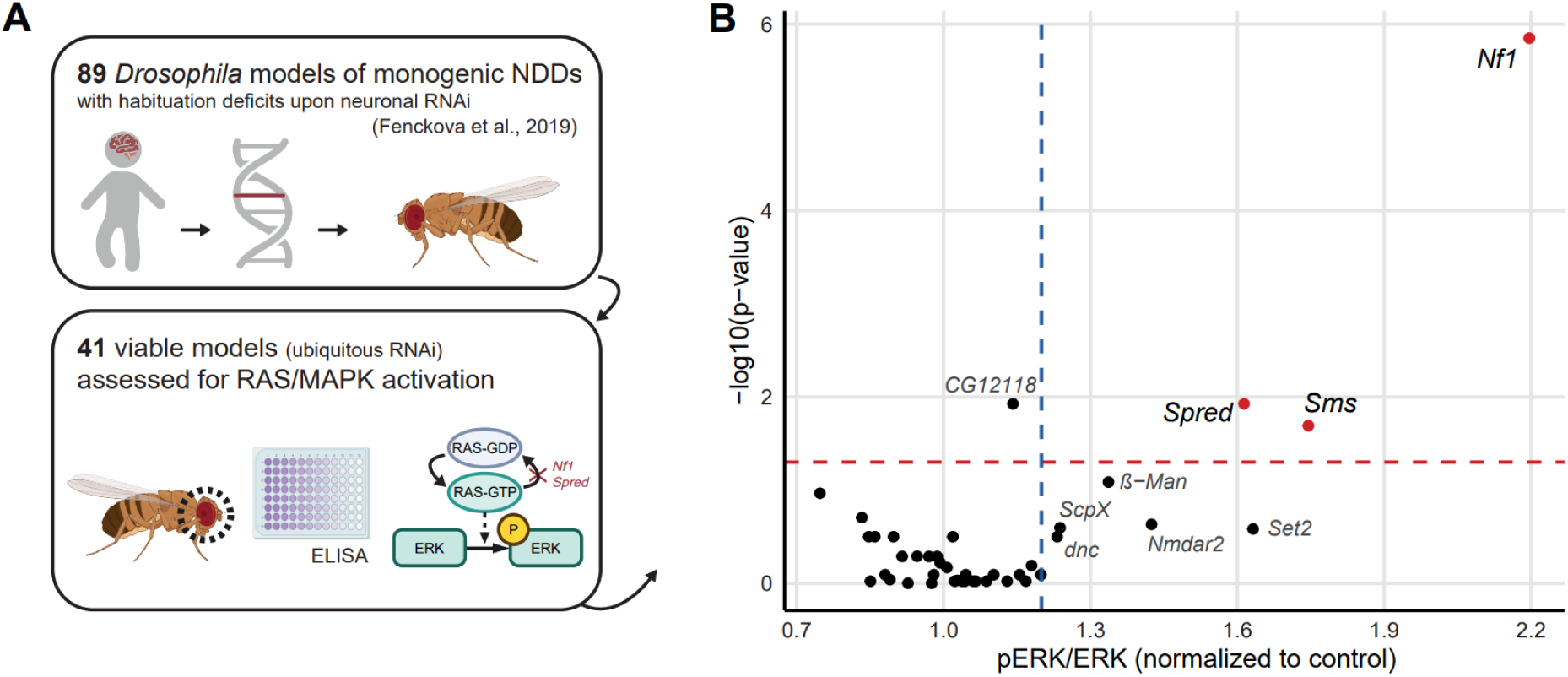
ELISA-based screen for RAS-MAPK activation in *Drosophila* NDD models with habituation deficits. **(A)** Schematic overview of the screening strategy. A total of 89 RNAi lines targeting NDD-associated genes and previously shown to cause habituation learning deficits when expressed pan-neuronally were selected. Ubiquitous knockdown (*Act-Gal4 > UAS-RNAi*) yielded 41 viable models. Head lysates were subjected to ELISA to quantify pERK and ERK levels; their ratio serves as a readout of RAS/MAPK activity. *Nf1* and *Spred* are highlighted as established negative regulators, whose loss-of-function increases RAS/MAPK signaling and thus elevates pERK/ERK ratios. **(B)** Results from the ELISA screen. Mean pERK/ERK fold change of knockdown models normalized to controls (*Act-Gal4 / control*) are plotted against the FDR-corrected p-values estimated using a linear model as described in Materials and Methods. Per genotype ≥3 biological replicates are included (except for *Set2*, n=1 due to lethality). Red line indicates p = 0.05, blue line indicates fold change > 1.2.

### Loss of *Sms* phenocopies *Nf1* with increased RAS/MAPK signaling

To validate the ELISA screen findings and further characterize *Sms* as a potential regulator of RAS/MAPK signaling, we performed western blot analysis on adult head extracts from independent *Sms* knockdown flies (*Act-Gal4 > Sms^RNAi^*). This confirmed a significant increase in the pERK/ERK ratio, consistent with RAS/MAPK pathway overactivation (Fig. 2A, **A’**). Interestingly, based on normalization with β-tubulin, this elevated pERK/ERK ratio may result from a reduction in ERK rather than an increase in pERK, as reflected by decreased ERK/β-tubulin but unaltered pERK/β-tubulin levels.

**Figure 2.**
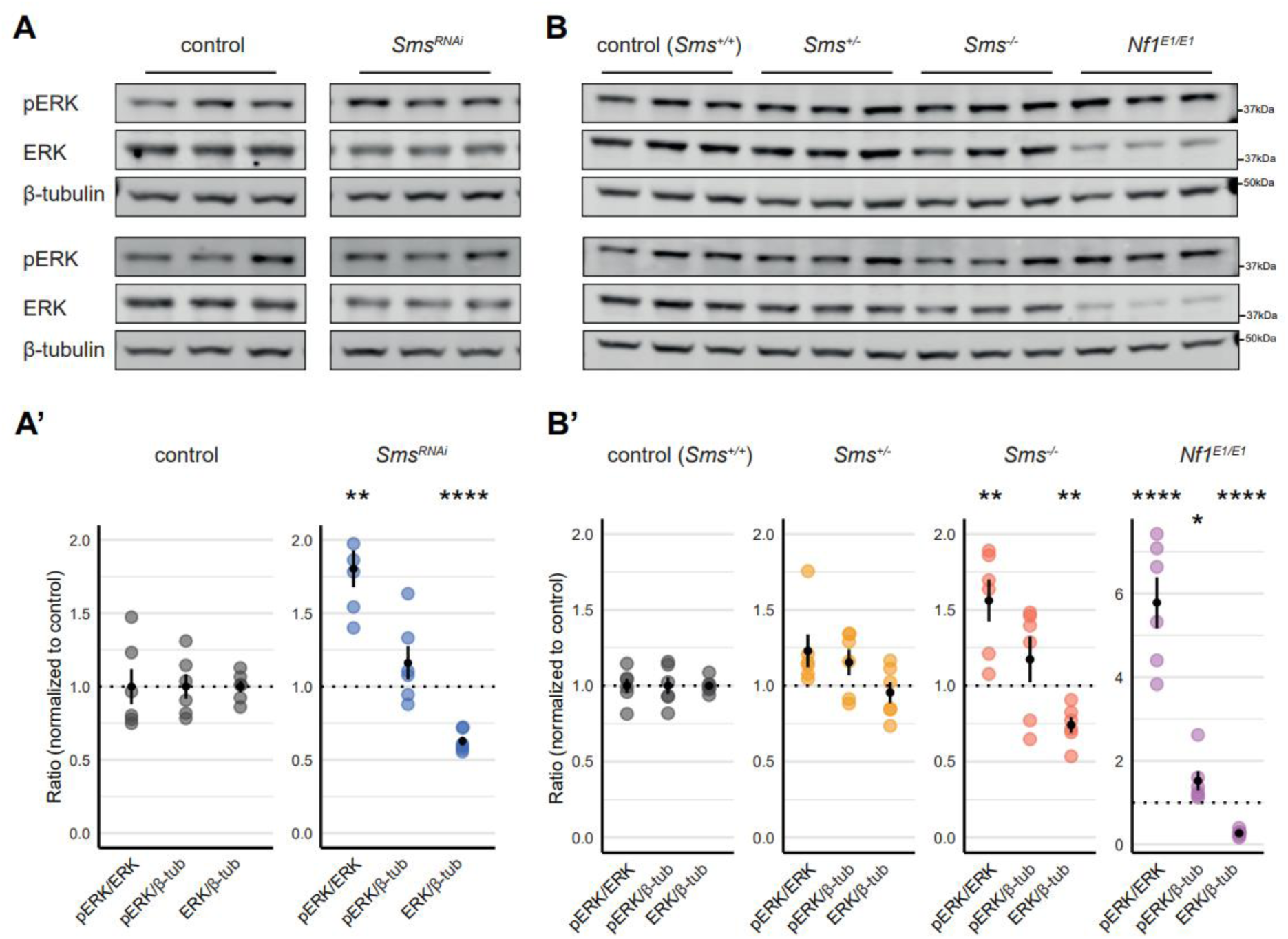
Loss-of-*Sms* dysregulates RAS-MAPK in a similar way as *Nf1*. **(A)** Western blots of head lysates from *Sms* knockdown models (*Act-Gal4 > Sms^RNAi^*) and controls (*Act-Gal4 / control*). **(A’)** Quantification and normalization to controls reveal that elevated pERK/ERK is driven by a decrease in ERK/β-tubulin (6 biological replicates, 2 blots). **(B)** Western blots of head lysates from *Sms* and *Nf1* complete loss-of-function mutants. **(B’)** Quantification and normalization to controls show the same pattern as *Sms* knockdown models (6 biological replicates, 2 blots). Graphs indicate mean ± SEM. Statistical significance was assessed using linear models as described in Materials and Methods. Corrected p-values are indicated as follows: * p < 0.05, ** p < 0.01, **** p < 0.0001.

To confirm these findings in an independent genetic loss-of-function model, we analyzed a previously characterized, viable *Sms* null mutant (*Sms^-/-^*) (Li et al., 2017). Western blots from *Sms^-/-^*head lysates recapitulated the increased pERK/ERK and decreased ERK/β-tubulin ratios observed in the knockdown animals (Fig. 2B, **B’**). *Sms* heterozygotes (*Sms^+/-^*) showed intermediate pERK/ERK and ERK/β-tubulin levels, illustrating a dose-dependent relationship between *Sms* levels and ERK homeostasis.

To facilitate interpretation of the observed ERK levels and activity, we included a classic RASopathy model into our analysis. *Nf1^E1/E1^* mutants, which carry homozygous alleles containing loss-of-function variants of *Nf1* generated through ethyl methanesulfonate (EMS) mutagenesis (Walker et al., 2006), displayed the same, even more pronounced pattern of increased pERK/ERK ratios in the presence of lower ERK levels (Fig. 2B, **B’**). The fact that loss of *Sms* recapitulates *Nf1* supports that *Sms* may act as a novel upstream modulator of ERK signaling.

### Loss of *Sms* causes hyperreactivity and habituation deficits

Building on these molecular findings, we next asked whether loss of *Sms* also recapitulates behavioral phenotypes previously associated with RAS/MAPK dysregulation. Because sensory processing alterations in individuals with RASopathies and their models encompass both baseline sensory reactivity and habituation learning (Carreno-Munoz et al., 2021; Pride et al., 2023; Wolman et al., 2014), our goal here was to systematically evaluate both components in *Sms* mutants. Prior work suggested that pan-neuronal *Sms* knockdown impairs habituation learning in the *Drosophila* light-off startle paradigm (Fenckova et al., 2019), together raising a possibly conserved role for *Sms* in sensory filtering in flies. To determine whether these effects extend to a genetic full loss-of-function model and to characterize the sensory profile, we assessed *Sms* mutants using two inter-trial intervals in the light-off startle paradigm (Fig. 3A). In this paradigm, flies jump in response to abrupt lights-off stimuli. When stimuli are presented at five-second intervals, flies do not habituate, and the resulting jump rate quantifies baseline sensory reactivity, though potentially masked by motor impairments (Fenckova et al., 2019; Stessman et al., 2017). In contrast, one-second interval stimulation induces a gradual decline in jump responses, reflecting habituation learning. Following suggested analytical approaches (McDiarmid et al., 2017), we quantified habituation by calculating a habituation ratio: the average jump rate during trials 51–100 (the habituated plateau) normalized to each fly’s unhabituated baseline reactivity measured under the five-second interval condition (Fig. 3A). This approach estimates the extent to which flies reduce their responses during the habituation assay, independent of differences in baseline reactivity.

**Figure 3.**
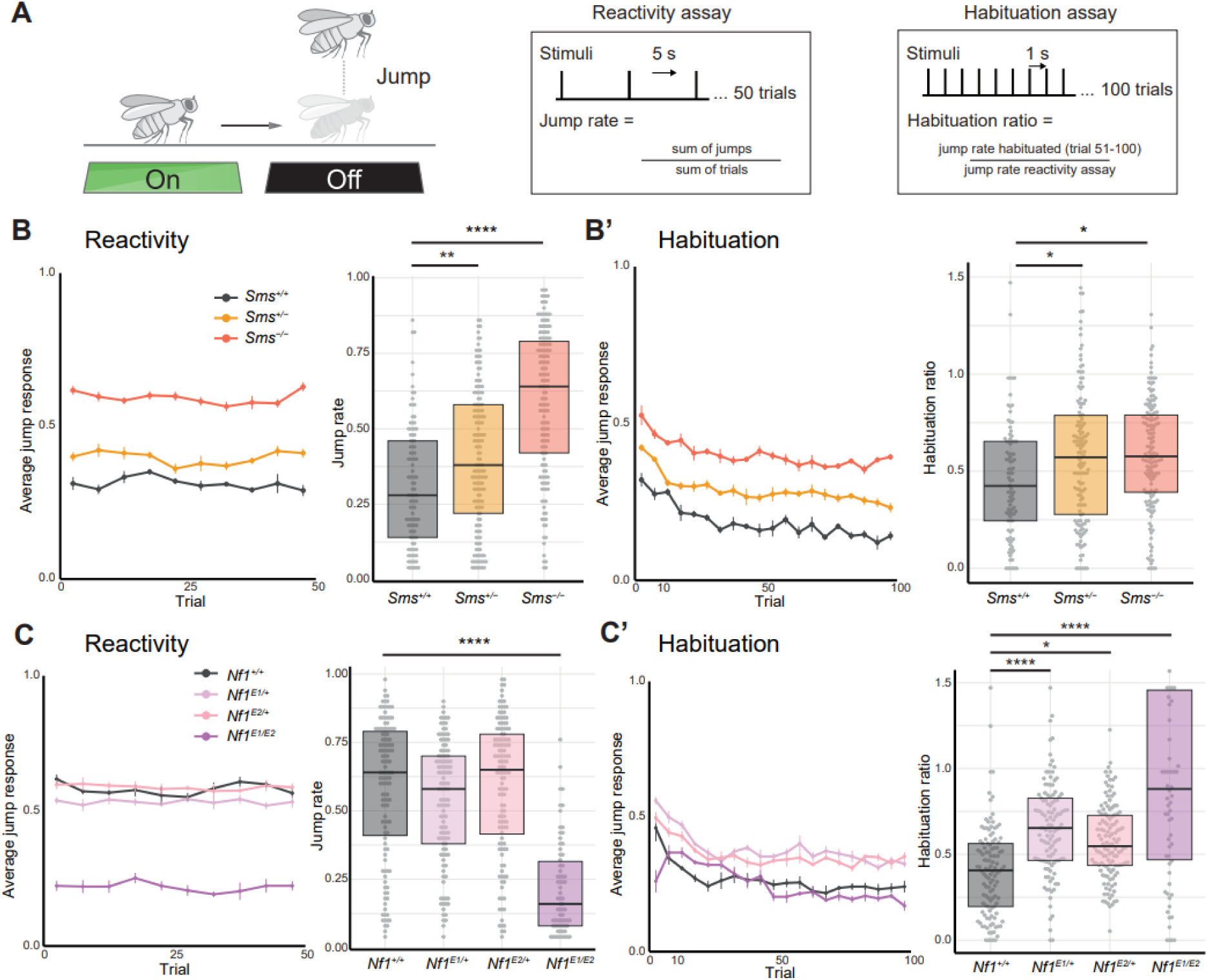
Loss of *Sms* leads to hyperreactivity and habituation deficits in the light-off startle paradigm. **(A)** In the light-off startle paradigm, flies jump in response to brief (15 ms) light-off stimuli. Reactivity was measured as jump rate under 5 s interval stimulation (reactivity assay). Habituation was quantified as the ratio of jump rate during trials 51–100 under 1 s interval stimulation to jump rate in the reactivity assay. **(B)** *Sms^-/-^* mutants show elevated jump rates in the reactivity assay compared to controls, with *Sms^+/-^* flies displaying intermediate phenotypes (4-5 replicates, 105-153 total flies per genotype). **(B’)** Habituation ratios are significantly elevated in *Sms^-/-^* and *Sms^+/-^* mutants, indicating impaired habituation even after accounting for increased baseline reactivity (4-5 replicates, 93-151 total flies per genotype). **(C)** *Nf1* heterozygous mutants have no altered baseline jump rates, but *Nf1^E1/E2^* flies show strongly reduced jump rates, potentially indicating motor impairment (4 replicates, 70-128 total flies per genotype). **(C’)** Habituation ratios are significantly elevated in hetero- and transheterozygous *Nf1* mutants, indicating impaired habituation (4 replicates, 66-126 total flies per genotype). Line graphs indicate mean ± SEM jump responses per bins of 5 stimuli. Boxes indicate median ± 25th and 75th percentiles. Statistical significance was assessed using linear models as described in Materials and Methods. p-values are indicated as follows: * p < 0.05, ** p < 0.01, **** p < 0.0001.

*Sms^-/-^* mutants displayed significantly increased jump rates in the reactivity assay compared to controls, with *Sms^+/-^* heterozygotes showing an intermediate, still significant phenotype (Fig. 3B). In the habituation assay, both *Sms^-/-^* and *Sms^+/-^* mutants exhibited elevated habituation ratios relative to controls (Fig. 3**B****’**). This indicates that even after accounting for their increased baseline reactivity, mutant flies maintained elevated reactivity during the habituated phase, suggesting impaired adaptive filtering.

To compare these findings with a classic RASopathy model, we again tested Nf1 mutant flies. Baseline reactivity of heterozygous loss-of-function (*Nf1^E1/+^* and *Nf1^E2/+^*) flies was unaffected, whereas transheterozygous *Nf1^E1/E2^* null mutants showed reduced baseline reactivity, indicative of motor impairments (Fig. 3C). Despite this limitation, all *Nf1* mutant conditions exhibited elevated jump responses in the habituated state when normalized to their baseline reactivity, revealing habituation deficits (Fig. 3**C****’**). While the hyperreactive phenotype of *Sms* mutants was not recapitulated in *Nf1* mutants, the shared habituation deficits could suggest a converging role in adaptive sensory processing.

### *Sat* knockdown mirrors *Sms*-associated sensory and RAS/MAPK phenotypes

*Sms* encodes spermine synthase, an enzyme that converts spermidine into spermine within the evolutionarily conserved polyamine pathway (Fig. 4A) (Burnette and Zartman, 2015; Wu and Liu, 2024). To determine whether the sensory processing phenotypes observed in *Sms*-deficient models reflect a broader consequence of polyamine pathway disruption, we ubiquitously knocked down and tested five additional pathway components. We included RNAi lines (RNAi-1) targeting *Sat*, *Odc1*, *Odc2*, *SamDC*, and *SpdS* from the VDRC GD library. We were able to include a second, independent RNAi line (RNAi-2) for *Sat*, *Odc2*, and *SpdS* from the VDRC KK library.

**Figure 4.**
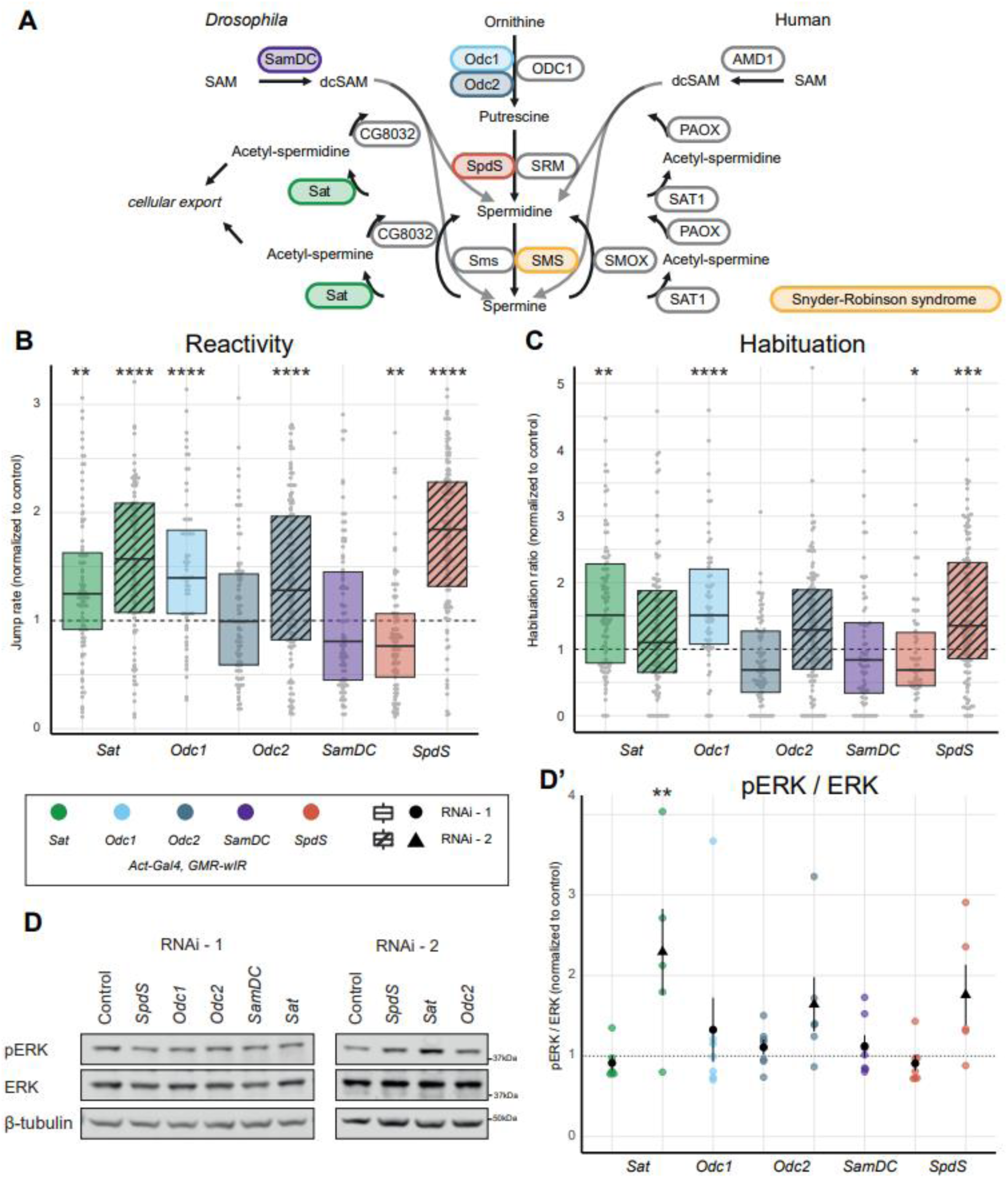
Knockdown of polyamine metabolism genes identifies sensory processing and RAS-MAPK phenotypes upon loss-of-*Sat*. **(A)** *Drosophila* (left) and human (right) genes mapped onto the enzymes involved in polyamine metabolism, based on Burnette and Zartman (2015); Wu and Liu (2024). **(B)** Jump rates in reactivity assay and **(C)** habituation ratios of knockdown models (*Act-Gal4,GMR-wIR > UAS-RNAi*) normalized to controls (*Act-Gal4,GMR-wIR / control*; 3-4 replicates, 62-128 total flies per genotype). Boxes indicate median ± 25th and 75th percentiles. **Fig. S1** for line graphs and non-normalized data. **(D)** Illustrative western blot bands and **(D’)** quantifications of pERK/ERK ratios in lysates of 10 pooled fly heads collected after behavioral testing (5-7 samples per genotype, **Fig. S2** for all blots and ratios). Graphs indicate mean ± SEM. Statistical significance was assessed using linear models as described in Materials and Methods. p-values are indicated as follows: * p < 0.05, ** p < 0.01, *** p < 0.001, **** p < 0.0001.

Knockdown of *Sat* (RNAi-1 and −2), *Odc1*, *Odc2*, and *SpdS* led to elevated jump responses in the reactivity assay, sharing increased sensory reactivity as phenotypes with *Sms* (Fig. 4B). Impaired habituation, as reflected by significantly elevated habituation ratios, was observed in single RNAi lines targeting *Sat*, *Odc1*, and *SpdS* (Fig. 4C). After being assayed for reactivity and habituation, fly heads from all genotypes were used for determining pERK, ERK and β-tubulin by western blot analysis. *Sat^RNAi-2^*showed a significant increase in pERK/ERK ratios (Fig. 4D, **D’**). The increase was driven by elevated pERK/β-tubulin levels, with no changes in ERK/β-tubulin (**Figs. 4D** and **S2**). This pattern is consistent with RAS/MAPK pathway overactivation, although it exhibits a distinct molecular signature compared to the decreased ERK/β-tubulin levels observed in *Sms-* and *Nf1*-deficient models.

Taken together, knockdown of *Sat* induces hyperreactivity across two independent RNAi lines, while habituation deficits and RAS/MAPK pathway overactivation are each observed in one seperate line. These findings highlight *Sat* as the only gene whose knockdown recapitulates all three phenotypes seen in *Sms*-deficient models. *Sat* encodes spermidine/spermine acetyltransferase, the rate-limiting enzyme in spermidine catabolism (Fig. 4A). The fact that *Sms* and *Sat* both use spermidine as a substrate to generate different products (spermine versus acetyl-spermidine) suggests that spermidine accumulation may underlie the convergent behavioral and molecular phenotypes.

### GABAergic origin of sensory processing deficits in *Sms* and *Sat* models

Given the ubiquitous role of polyamines (Sagar et al., 2021), we next sought to determine whether the sensory phenotypes observed in *Sms* and *Sat* models arise from a shared cellular context. Previous work has shown that pan-neuronal knockdown of *Sms* impairs habituation (Fenckova et al., 2019). We therefore first used *Elav-Gal4* to drive pan-neuronal RNAi-mediated knockdown of both enzymes. Unlike seen in the *Sms* mutant, pan-neuronal knockdown of *Sms* does not significantly increase baseline reactivity (Fig. 5A). However, consistent with findings from Fenckova et al. and *Sms* mutant flies, the habituation ratio is significantly increased (Fig. 5B). Looking at the response curves, both *Sat* RNAi lines exhibited elevated jump responses during the habituated phase (Fig. 5B). However, because these lines also showed increased baseline reactivity (Fig. 5A), the habituation ratio remained unchanged, suggesting that deficits in habituation may be secondary to a heightened sensory reactivity.

**Figure 5.**
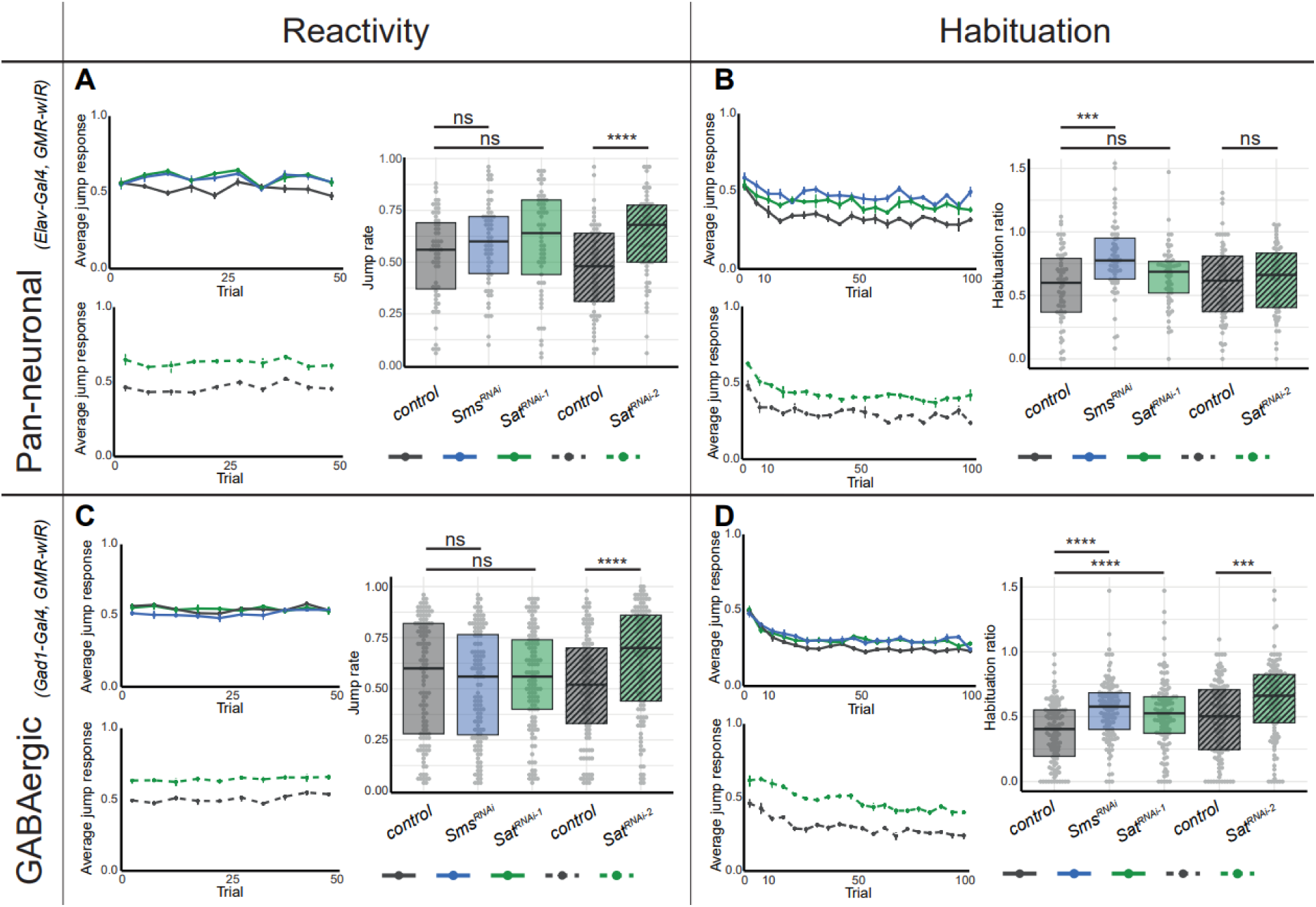
*Sms* and *Sat* cause habituation deficit when knocked down specifically in GABAergic neurons. **(A)** Jump rates in reactivity assay and **(B)** habituation ratios of pan-neuronal knockdown models (*Elav-Gal4, GMR-wIR > UAS-RNAi)* normalized to controls (*Elav-Gal4,GMR-wIR / control)* reveal significantly elevated reactivity in *Sat^RNAi-2^* and habituation ratios in *Sms^RNAi^* (4 replicates, 116-124 total flies per genotype). **(C)** Jump rates in reactivity assay and **(D)** habituation ratios of GABAergic knockdown models (*Gad1-Gal4, GMR-wIR > UAS-RNAi)* normalized to controls (*Gad1-Gal4,GMR-wIR / control)* reveal significantly elevated reactivity in *Sat^RNAi-2^* and habituation ratios in *Sms^RNAi^*, *Sat^RNAi-1^* and *Sat^RNAi-2^* (2 replicates, 61-63 total flies per genotype). Line graphs indicate mean ± SEM jump responses per bins of 5 stimuli. Boxes indicate median ± 25th and 75th percentiles. Statistical significance was assessed using linear models as described in Materials and Methods. p-values are indicated as follows: *** p < 0.001, **** p < 0.0001.

Habituation deficits in RASopathy models, including *Nf1*, have previously been attributed to dysfunction within GABAergic neurons (Fenckova et al., 2019). To test whether perturbation of polyamine metabolism in these neurons is also sufficient to induce cognitive phenotypes, we selectively knocked down *Sms* and *Sat* using the GABAergic driver *Gad1-Gal4*. Baseline reactivity remained unchanged for *Sms^RNAi^* and *Sat^RNAi-1^*, whereas *Sat^RNAi-2^* showed a significant increase (Fig. 5C). Notably, despite elevated baseline reactivity under GABAergic-specific knockdown, *Sat^RNAi-2^* displayed habituation deficits (Fig. 5D), unlike when crossed to ubiquitous (Fig. 4C) or pan-neuronal (Fig. 5B) drivers. Moreover, increased habituation ratios were also observed in *Sms^RNAi^* and *Sat^RNAi-1^*, indicating that all three RNAi lines exhibited impaired habituation when crossed to the GABAergic driver (Fig. 5D). Together, this identifies an essential role for both *Sms* and *Sat* in GABAergic neurons for sensory processing, with *Sms* appearing to be primarily involved in habituation learning, while *Sat* contributes to both increased sensory reactivity and impaired habituation. This suggests that knockdown models of polyamine metabolism genes can recapitulate the cellular origin of habituation deficits in classic RASopathy models.

## DISCUSSION

RASopathies are developmental conditions marked by elevated RAS/MAPK signaling, a pathway previously identified as a central node in impaired habituation learning across *Drosophila* models of ID (Fenckova et al., 2019). We reasoned that some of the habituation deficit disease models may represent “hidden RASopathies”: genetic diseases not classically linked to RAS/MAPK signaling but sharing molecular and clinical features. Here, we report several disorders whose *Drosophila* model, in addition to habituation deficits, exhibits RAS/MAPK overactivation. Notably, this screen highlighted polyamine metabolism as a modulator of RAS/MAPK activity. Manipulating additional polyamine pathway enzymes further demonstrated a direct influence on both RAS/MAPK activity and sensory processing behavior, with perturbation in inhibitory neurons alone being sufficient to recapitulate the phenotypes. These findings extend the molecular framework underlying habituation deficits and position polyamine metabolism as a novel regulator of RAS/MAPK-driven sensory processing, with implications for genetic disorders linked to RAS and polyamine pathways.

### Relevance of the screening approach for rare genetic neurodevelopmental disorders

Our screening approach was designed to measure RAS/MAPK activity in fly heads of habituation-deficient models, providing disease-relevant resolution. Using this strategy, we identified modifiers, including most notably *Sms*, that were not detected in previous *in vitro* RAS/MAPK screens (Ashton-Beaucage et al., 2014; Sawyer et al., 2020). We report three genes, *Nf1*, *Spred* and *Sms*, whose knockdown produced pERK/ERK ratios above the significance threshold. *Sms* showed an intermediate pERK/ERK level relative to the classic RASopathy genes *Nf1* and *Spred*, supporting its identification as a genuine RAS/MAPK modulator. *Sms* encodes spermine synthase, an enzyme involved in polyamine metabolism. In humans, pathogenic variants in *SMS* cause Snyder-Robinson syndrome, an X-linked NDD characterized by ID, muscle hypotonia, and skeletal abnormalities (Cason et al., 2003; Snyder and Robinson, 1969). In addition, five further genes, *Set2* (*SETD2* in human), *Nmdar2* (*GRIN2A/B*), *β-Man* (*MANBA*), *ScpX* (*SCP2*), and *dnc* (*PDE4A-D*), showed more than 1.2-fold increases in pERK/ERK levels.

Notably, two of these genes, *Nmdar2* and *dnc*, have known mechanistic links to RAS/MAPK signaling, reinforcing their relevance even below statistical cutoffs. *Nmdar2* encodes an NMDA receptor subunit essential for synaptic plasticity. NMDA activation can both stimulate Ras and limit its activity via SynGAP1, a RASopathy-associated protein (Jeyabalan and Clement, 2016; Kim et al., 2005; Wang et al., 2007). *Dnc* encodes the fly orthologue of mammalian PDE4 family of phosphodiesterases (PDE4s), which degrade cAMP, a second messenger that modulates signaling pathways including RAS/MAPK (Donders et al., 2024). Further support for a role of cAMP in the RAS-associated habituation phenotype comes from the fact that *Nf1* can directly control both cAMP and RAS/MAPK pathways (Botero and Tomchik, 2024), and inhibition of PDE4 improved habituation in NF1 zebrafish models (Wolman et al., 2014). Considering these established mechanistic links, the increase in RAS/MAPK activation observed for these genes – albeit non-significant – may reflect subtle or potentially context-dependent contributions to pathway regulation. Our identification of these associations in genetic *in vivo* disease models warrants further investigation.

*Sms* knockdown increased pERK/ERK ratios, driven by reduced ERK/β-tubulin, a pattern consistent with RAS/MAPK overactivation as it is resembling the phenotype observed in *Nf1* mutants. Although prior work on the same *Nf1* allele reported elevated pERK with stable ERK and β-tubulin levels in adult head samples (Walker et al., 2006), this difference may reflect time- and context-dependent negative feedback on ERK expression after sustained pathway activation (Lake et al., 2016). One potential mechanism for such feedback involves pERK stimulated, RREB1-dependent induction of miR-143/145, which has been shown to attenuate ERK expression (Kent et al., 2013). Eventually, the balance between pERK and ERK is evidently disturbed, with predictably detrimental consequences, given the pathway’s reliance on tight spatiotemporal control (Ram et al., 2023).

### Spermidine as a regulator of RAS/MAPK signaling

We investigated additional components of the polyamine pathway to further establish their connection to sensory processing and RAS/MAPK dysregulation. In addition to *Sms*, genetic knockdown of *Sat* (*SAT1, SAT2, SATL1* in human) increased RAS/MAPK signaling, strengthening the link between this pathway and polyamines—molecules which are present across all cell types and living organisms, essential for nucleic acid stabilization, protein translation, and signal transduction (Xuan et al., 2023). Since both *Sms* and *Sat* use spermidine (SPD) as a substrate, loss of either gene product is expected to cause SPD accumulation, providing a possible pathogenic mechanism for the observed phenotypes. Interestingly, increased levels of polyamines are a hallmark of cancer (Sagar et al., 2021), and increased levels of spermidine specifically have been associated with elevated pERK in cancer cell cultures (Bachrach et al., 2001). Pharmacological reduction of polyamine synthesis via ODC inhibition (DMFO) has shown to be an effective cancer treatment (Schramm et al., 2025). Conversely, while DMFO is effective in reducing putrescine (PUT) and SPD levels, spermine (SPM) levels go up (Flamigni et al., 1999). Indeed, SPM is proposed to have negative regulating effects on RAS/MAPK as evidenced from its capability to disturb phosphorylation of DRaf (Stark et al., 2011).

In addition, our findings propose that increased RAS/MAPK signaling may contribute to cognitive impairments in Snyder-Robinson syndrome, caused by pathogenic variants in *SMS*. In fly models of SRS, elevated SPD has been linked to cellular toxicity through ROS-generating acetylation and oxidation (Li et al., 2017). Notably, recent work in human bone-marrow–derived pluripotent stromal cells demonstrated that upregulated SAT1 strongly reduced SPD levels (Cressman et al., 2024). This aligns with our observation that both *Sms* loss and *Sat* loss lead to phenotypes associated with SPD accumulation. The *Sms^G56S^* mutant mouse model exhibits mitochondrial respiration defects and increased energy expenditure, along with abnormal growth and body composition, and behavioral phenotypes such as an increased and maintained fear response (Akinyele et al., 2024). Upon reanalysis of the RNA-seq data from this mouse model, we identified an enrichment of dysregulated genes within RAS- and ERK-related Gene Ontology categories, including “Ras protein signal transduction” (GO:0007265) and “ERK1 and ERK2 cascade” (GO:0070371) (**Table S3**). This supports our finding of dysregulated RAS/MAPK signaling in a model of SRS and extends it across species. While polyamine imbalance could drive RAS/MAPK activation, the pathway may also feed back onto polyamine metabolism. In cancer cells, inhibition of MEK (PD98059) can suppress expression of *ODC1* and thus restore polyamine balance (Flamigni et al., 1999), suggesting that restoring RAS/MAPK holds therapeutic potential to restore polyamine balance in SRS.

### Polyamine dysregulation in GABAergic neurons drives habituation deficits

Behavioral assays revealed that *Sms* and *Sat* mutants display impairments in both baseline sensory reactivity and habituation learning, as assessed using the light-off startle paradigm. Although habituation and sensory reactivity represent distinct aspects of sensory processing, namely adaptation to repeated stimuli versus general responsiveness, they are both fundamental components of how organisms respond to sensory input (He et al., 2023; McDiarmid et al., 2017; Schauder and Bennetto, 2016). Importantly, both traits are clinically relevant. Altered sensory responsiveness and habituation are frequently observed in individuals with neurodevelopmental disorders and RASopathies, as well as in their animal models (Carreno-Munoz et al., 2021; Ethridge et al., 2019; Pride et al., 2023; Wolman et al., 2014). Their relative importance to atypical behavioral responses and problems in daily living skills is however unknown.

Neuronal subtype-specific knockdown experiments demonstrated that both *Sms* and *Sat* operate in GABAergic neurons, pinpointing inhibitory circuitry as the origin of dysfunction. Polyamines intersect with inhibitory transmission via putrescine, a precursor for GABA (Makletsova et al., 2022). While increased putrescine availability can enhance GABA synthesis and tonic inhibition in epilepsy models (Kovács et al., 2021), chronic accumulation of spermidine and putrescine in *Sms* and *Sat* mutants may destabilize inhibitory balance, paralleling findings in NF1 models where enhanced GABA release contributes to cognitive deficits (Costa et al., 2002; Cui et al., 2008; Omrani et al., 2015). Furthermore, it aligns with observed habituation defects when manipulating RASopathy genes, including *Nf1* in GABAergic neurons (Fenckova et al., 2019). Together, these results suggest that polyamine dysregulation and RAS/MAPK hyperactivation converge on inhibitory circuit dysfunction, producing deficits in sensory reactivity and habituation learning.

### Shared pathophysiology of polyaminopathies and RASopathies creates therapeutic opportunities

The convergence of polyaminopathies and RASopathies extends beyond molecular and behavioral phenotypes. Snyder-Robinson syndrome shares clinical features with NF1, Noonan, and Costello syndromes, including cognitive/learning impairments, but also short stature, reduced bone mineral density, and skeletal abnormalities such as scoliosis (Aftab and Dattani, 2019; Friedman, 1993; Gripp et al., 2019; Rauen and Tidyman, 2024; Reynolds et al., 2025), further supporting shared aspects of pathophysiology. This overlap has important therapeutic implications, as inhibitors of MEK are FDA-approved for tumors in NF1 (Cook, 2025; Lalancette et al., 2024; Walsh et al., 2021). At the same time, polyamines can be targeted by ODC inhibitor Eflornithine (DFMO), which was developed to treat cancer and repurposed for Bachmann-Bupp syndrome (OMIM #619075), caused by activating variants in *ODC1* (Bachmann et al., 2024). Eflornithine is also proposed as a treatment in SRS, as it restored SPD:SPM balance and improved several readouts in human bone marrow-derived pluripotent stromal cells and longevity in fly models (Cressman et al., 2024; Stewart et al., 2023).

By highlighting shared signaling disruptions across genetically distinct but functionally convergent rare neurodevelopmental disorders, these findings raise the potential for unified therapeutic strategies targeting both polyamine metabolism and RAS/MAPK signaling.

## METHODS

### *Drosophila* stock selection for ELISA screen

146 genes, when pan-neuronally knocked down, have been associated with habituation defects in our light-off startle paradigm. Before screening, 57 genes were excluded because the gene was either not associated with monogenic NDD in the SysNDD database curated at 12-nov-2025 (13 genes), the habituation deficit fly line was from another source than the Vienna *Drosophila* Resource Center (VDRC) GD or KK collection (6 genes), or was not readily available (2 genes), the UAS-RNAi construct was located on the X-chromosome (13 genes) or at the 40D landing-site (23 genes). All excluded RNAi lines are listed in **Table S1**, and included RNAi lines are listed in **Table S2** including a reference to the study where a habituation deficit is shown with a pan-neuronal driver, if published.

### *Drosophila* stocks and maintenance

Fly stocks were maintained on standard cornmeal-yeast-sugar-agar medium supplemented with the mold inhibitors methyl paraben and propanoic acid, at 25 °C and 60% humidity under a 12:12 h light-dark cycle. Genotypes are referred to by shorthand labels in figures and text, with full genotypes provided in **Table S4**. Gene knockdown was achieved using the Gal4/UAS system (Brand and Perrimon, 1993). Knockdown animals were generated by crossing Gal4 driver lines to UAS-RNAi lines from the GD and KK libraries of VDRC (Dietzl et al., 2007) **(Table S1 and S4)**. Control animals were generated by crossing each Gal4 driver line to the genetic background control lines of the corresponding library (VDRC #60000 for GD lines and #60100 for KK lines). The *Sms* mutant line (**Table S4**) has a P-element insertion resulting in less than 0.05% transcript levels of *Sms* (Li et al., 2017) and was backcrossed 7 times to the *w-* control line (VDRC #60000), which served as control. EMS generated *Nf1^E1^* and *NF1^E2^*mutant and isogenic control lines were kindly provided by James Walker (Walker et al., 2006). In several models (**Table S4**), a GMR-wIR element (eye-specific RNAi knockdown of *white*) was introduced to suppress eye pigmentation which is required for light-off startle responses (Fenckova et al., 2019).

### Enzyme-Linked Immunosorbent Assay (ELISA)

Phospho-ERK (pERK) and ERK levels were quantified in fly head extracts using the ERK1/2 (pT202/Y204) + Total ERK1/2 ELISA kit (Abcam, ab176640). Twenty 1–3-day-old flies were anesthetized with CO₂, snap-frozen in liquid nitrogen, and decapitated by flicking. Heads were collected on ice using a brush. Heads were homogenized in 100 µL of the provided lysis buffer using a plastic pestle, incubated on ice for 20 minutes, and centrifuged at 12,000 rpm for 20 minutes at 4 °C. Supernatants were diluted 1:10 to approximately ∼500 µg/mL protein. A positive control was prepared per kit instructions. For each well, 50 µL of diluted sample or control and 50 µL of antibody cocktail were added. Plates were incubated for 1 hour at room temperature (400 rpm), washed 3× with provided wash buffer, then incubated with 100 µL TMB substrate for 15 minutes in the dark. Reactions were stopped with 100 µL stop solution, and absorbance was read at 450 nm.

Samples were run in technical duplicates for both pERK and ERK and their respective mean absorbance was included only if the coefficient of variation (CV, calculated as standard deviation / mean) was ≤0.2 for both pERK and ERK. For each RNAi model ≥3 biological replicates (except for *Set2*, n=1 due to lethality) were tested, meaning independent crosses and plates, but always measured on the same plate together with a background control.

Statistical significance was assessed using a linear mixed-effects model (lme4 package) in R (v4.3.3) (Bates et al., 2015). Log₁₀-transformed pERK/ERK absorbance ratios were used as the response variable, with genotype as a fixed effect and ELISA plate as a random effect. Pairwise comparisons between disease models and their respective controls were performed using the emmeans R package (DOI: 10.32614/CRAN.package.emmeans), with multiple testing correction using the Benjamini-Hochberg false discovery rate (FDR). Data figures are generated using the ggplot2 R package (Wickham, 2016).

### Western blotting

To quantify pERK, ERK, and β-tubulin levels, western blotting was performed. For validation of *Sms* phenotypes (Fig. 2), heads from ten 1–3-day-old flies were collected as described for ELISA. For polyamine pathway knockdown models (Fig. 4), heads from ten 7–10-day-old flies were collected following light-off startle assays, anesthetized on ice, and snap-frozen in liquid nitrogen. Heads were homogenized in 60µL RIPA buffer with protease and phosphatase inhibitors, incubated on ice for 20 minutes, and centrifuged at 12,000 rpm for 20 minutes at 4 °C. 15µL of supernatant was mixed with 5µL sample buffer (100 µM DTT in NuPAGE LDS Sample Buffer, Invitrogen) and heated at 95℃ for 5 minutes. Samples (12 µL) or 3 µL of Odyssey molecular weight marker were loaded onto 4–12% Bis-Tris gels (NuPAGE 15 wells, Invitrogen) and electrophoresed in MOPS buffer (NuPage MOPS SDS Running Buffer, Invitrogen) at 120 V for ∼2 hours. Proteins were transferred to nitrocellulose membranes (Trans-Blot Turbo Transfer Pack, Bio-Rad) using the Trans-Blot Turbo system (Bio-Rad) at 25V for 7 minutes. Membranes were blocked for one hour in 10% bovine serum albumin (BSA, Sigma-Aldrich, A8806-1G), and then incubated overnight at 4°C in primary antibodies diluted in a 1:1 mix of 10% BSA and tris buffered saline buffer with 0.2% tween (TBST). The antibody against pERK (M8159, Sigma) was diluted 1:2500, and total ERK (9102S, Cell signaling) was diluted 1:500. After three 5-minute washes in TBST, membranes were incubated with secondary antibodies (goat anti-mouse, Alexa Fluor 680, A-21057, Thermo Fisher Scientific and goat anti-rabbit, IRDye 800, LI-COR, each 1:10000) for 1 hour at room temperature in the dark. Membranes were then washed, rinsed in TBS, and imaged on an Odyssey Infrared Scanner (Odyssey® DLx Imaging System, Li-cor). Membranes were reprobed with anti-β-tubulin (dilution 1:10000, AB_2315513, DSHB) for 1 hour at room temperature, followed by secondary antibody incubation and imaging as above.

Quantification was conducted using Image Studio software (version 4.0.21). Local background was subtracted to obtain signal intensity of each band. Genotypes are always compared against control samples on the same gel. Samples are always from independent biological replicates. Statistical significance was assessed using a linear mixed-effects model (lme4 package) in R (v4.3.3) (Bates et al., 2015). Log₁₀-transformed pERK/ERK, pERK/β-tubulin, or ERK/β-tubulin ratios were used as the response variable, with genotype as a fixed effect and gel as a random effect. Pairwise comparisons between groups were performed using the emmeans R package (DOI: 10.32614/CRAN.package.emmeans), with multiple testing correction via Šidák method. Data figures are generated using the ggplot2 R package (Wickham, 2016).

### Light-off startle paradigm

Sensory processing phenotypes, reactivity and habituation, were assessed using the light-off startle paradigm (Aktogen Ltd.) as previous described (Fenckova et al., 2019). In each round, 32 flies aged 7–10 days were individually placed in tubes. Following a 5-minute acclimatization period, flies were subjected to the habituation paradigm, receiving 100 light-off stimuli (15 ms) with a 1-second intertrial interval (ITI). Then, following a 2-minutes rest period the reactivity assay was started where 50 stimuli were delivered with a 5-second ITI. Wing vibrations during jump responses were recorded by microphones at both ends, and a jump was concluded when a threshold of 0.8 V was exceeded. Data were collected using custom LabVIEW software (National Instruments, Austin, TX). Reactivity was quantified for each fly as the proportion of trials in which a jump occurred out of the 50 trials, including only flies that responded more than once. Habituation was quantified for each fly as the jump rate during the presumed habituated phase (trials 51–100 of the habituation assay), normalized to that fly’s jump rate in the reactivity assay, and included only flies with more than one response in both assays.

Statistical significance was assessed using a linear mixed-effects model (lme4 package) in R (v4.3.3) (Bates et al., 2015). Reactivity or habituation ratios were used as the response variable in separate linear models, with genotype as a fixed effect and light-off chamber identity and date of testing as random effects. Pairwise comparisons between groups were performed using the emmeans R package (DOI: 10.32614/CRAN.package.emmeans), with multiple testing correction via Šidák method. Data figures are generated using the ggplot2 R package (Wickham, 2016).

### Re-analysis of SMS mouse model transcriptome data

Processed RNA-seq data for the *Sms^G56S^* mouse model, which carries a missense variant in the *Sms* gene, and wild-type controls were obtained from the original study (GSE226413) (Akinyele et al., 2024). Differentially expressed genes (DEGs) were identified using the following criteria: false discovery rate (FDR) < 0.05 and absolute log2 fold change > 0.5 or < −0.5. This thresholding yielded 4,760 DEGs. Gene ontology (GO) biological process (BP) enrichment analysis was performed to identify overrepresented functional categories. Statistical comparison between groups of genes was conducted using the BinfTools R package (https://github.com/kevincjnixon/BinfTools), specifically employing the GO_GEM function. The background set for enrichment analysis consisted of all annotated mouse genes. Significant GO terms related to “RAS protein signal transduction” and the “ERK1 and ERK2 cascade,” along with their corresponding up- and down-regulated genes (log2 fold change > 0.5 or < −0.5, respectively), are provided in **Table S3**.

## Acknowledgements

We thank James Walker, the VDRC and the Bloomington *Drosophila* Stock Center for providing *Drosophila* strains. We are grateful to F. Kampshoff, S. Letteboer, M. Aslanyan and S. Beersum for experimental or logistic support, and to all members of the Schenck lab for helpful discussion.

## Competing interests

The authors declare that they have no competing interests.

## Funding

This work was in part supported by a Vici grant from the Netherlands Organization for Health Research and Development (ZonMw, 09150181910022) to A.S, and a Radboud Excellence Fellowship to S.G.J.

## Data and resource availability

All relevant data and resources can be found within the article and its supplementary information.

## Author contributions statement

Conceptualization: B.v.R., S.G.J., A.S. Methodology: B.v.R.

Software: B.v.R., M.Bo., S.G.J. Formal analysis: B.v.R., S.G.J.

Investigation: B.v.R., M.d.W., K.P., Z.-A.G., P.S., M.Be., S.G.J.

Data curation: B.v.R., S.G.J. Visualization: B.v.R.

Supervision: B.v.R., S.G.J., A.S. Project administration: B.v.R., A.S. Funding acquisition: S.G.J., A.S.

Writing – original draft: B.v.R., S.G.J., A.S. Writing – review & editing: All authors

---

**Figure S1.**
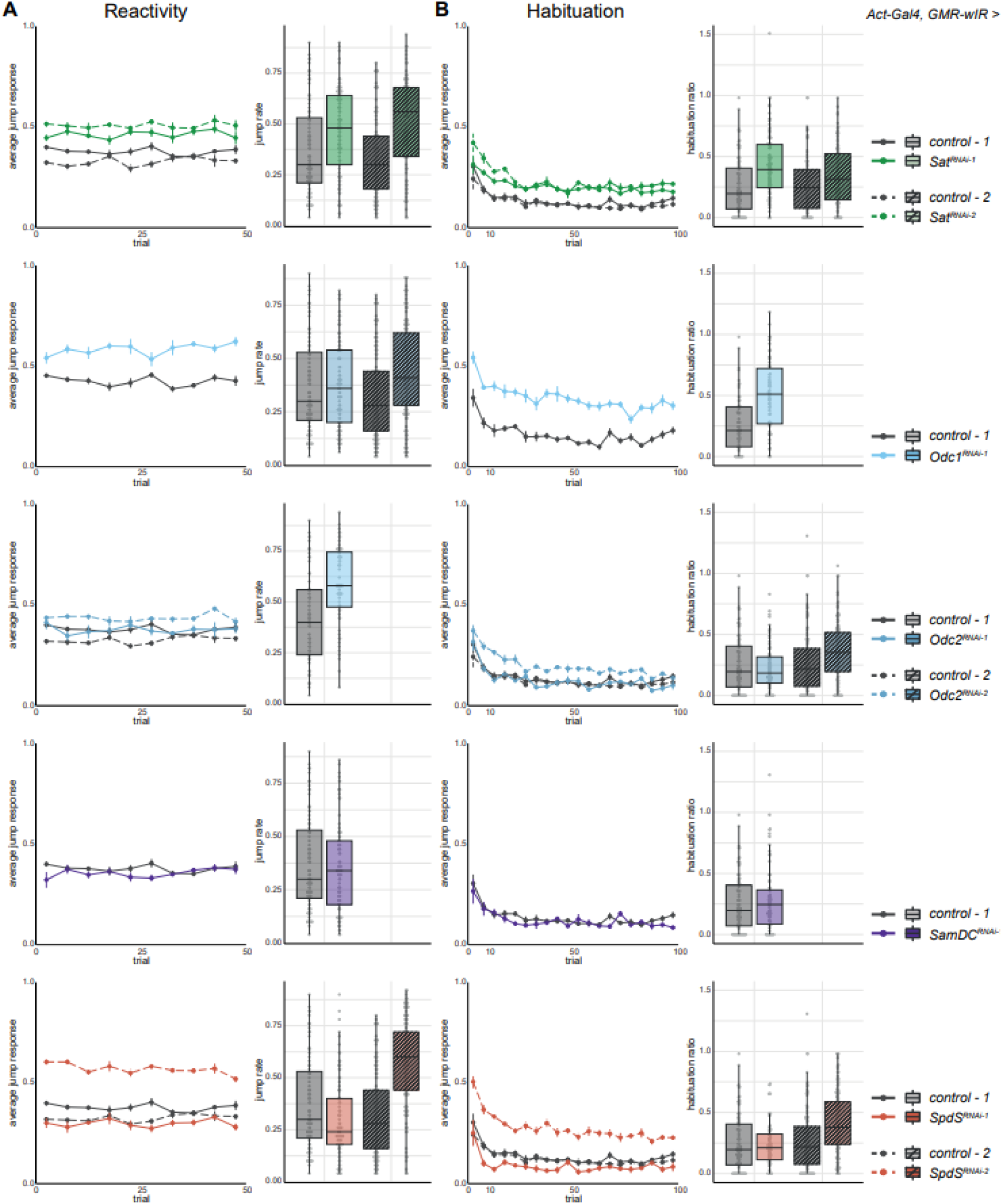
Sensory processing profiles upon knockdown of polyamine metabolism genes, related to Figure 4. (A) Jump rates in reactivity assay and (B) habituation ratios of knockdown models (*Act-Gal4,GMR-wIR > UAS-RNAi*) and controls (*Act-Gal4,GMR-wIR / control*; 3-4 replicates, 62-128 total flies per genotype). Line graphs indicate mean ± SEM jump responses per bins of 5 stimuli. Boxes indicate median ± 25th and 75th percentiles.

**Figure S2.**
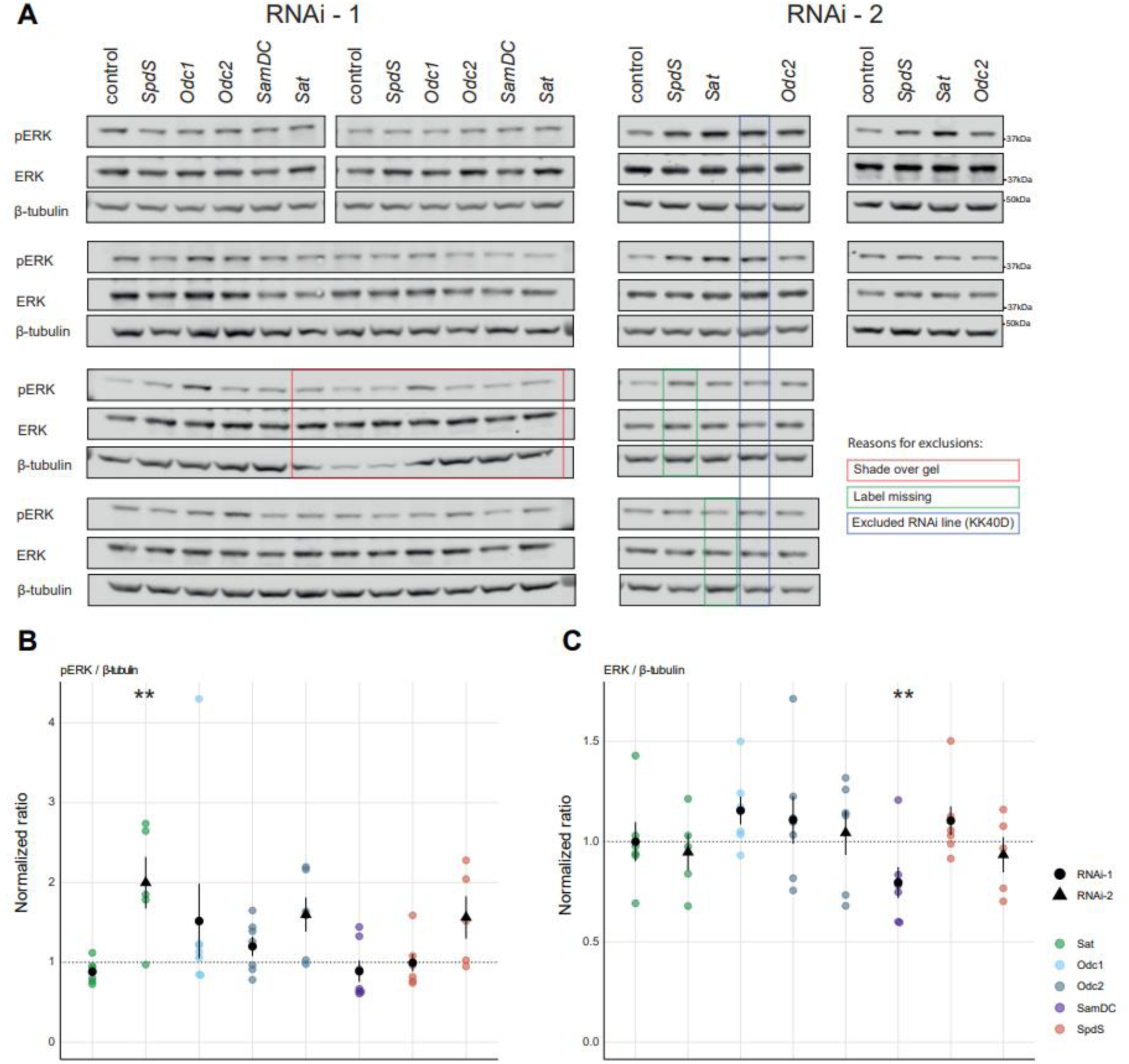
All western blots and ratios evaluating pERK, ERK and β-tubulin profiles upon knockdown of polyamine metabolism genes, related to Figure 4. (A) All western blot bands (labeled bands are excluded from quantification) and (B) quantifications of pERK/β-tubulin and (C) ERK/β-tubulin ratios in lysates of 10 pooled fly heads collected after behavioral testing (5-7 samples per genotype). Graphs indicate mean ± SEM.

## Supplementary table titles

**Table S1.** Results of ELISA screen on habituation deficit fly lines, related to Figure 1.

**Table S2.** List of excluded habituation deficit fly lines.

**Table S3.** RAS and ERK-related GO-terms enriched for differentially expressed genes in SmsG56S mouse model.

**Table S4.** List of fly lines used in this study.

## Notes

### Competing Interest Statement

The authors have declared no competing interest.

